# The unconventional Myosin-1C augments endothelial secretion of von Willebrand factor by linking contractile actomyosin machinery to the plasma membrane

**DOI:** 10.1101/2023.08.11.552954

**Authors:** Sammy El-Mansi, Tom P. Mitchell, Pika Miklavc, Manfred Frick, Thomas D. Nightingale

## Abstract

Blood endothelial cells control the hemostatic and inflammatory response by secreting von Willebrand factor (VWF) and P-selectin from storage organelles called Weibel-Palade bodies (WPB). Actin-associated motor proteins regulate this secretory pathway at multiple points. Prior to fusion, myosin Va forms a complex that anchors WPBs to peripheral actin structures allowing maturation of content. Post-fusion, an actomyosin ring/coat is recruited and compresses to forcibly expel the largest VWF multimers. Here we provide the first evidence for the involvement of class I myosins during regulated VWF secretion. We show that unconventional myosin-1C (Myo1c) is recruited post fusion via its pleckstrin homology domain in an actin-independent process providing a link between the actin ring and phosphatidylinositol 4,5-bisphosphate (PIP2) at the membrane of the fused organelle. This is necessary to ensure maximal VWF secretion in response to secretagogue stimulation. Inhibition of class I myosins using the inhibitor Pentachloropseudilin alters the kinetics of the exocytic actin ring. These data offer new insight into the control of an essential physiological process and provide a new potential way in which it might be therapeutically controlled.

**SIGNFICANCE STATEMENT:** Myosin motors play diverse roles in regulated secretion. In endothelial cells, the role of conventional myosins (e.g. non-muscle myosin II) are well described however little is known about the requirement of unconventional myosins. Our data identify an important function of the class 1 myosin, Myosin-1C, in the actomyosin mediated expulsion of an essential blood clotting factor (von Willebrand factor) from endothelial cells. This is the first description of how class 1 myosins contribute to primary hemostasis and is therefore greatly improves our understanding of a fundamental physiological process.

## MAIN TEXT

Endothelial cells (EC) contain rod-shaped storage organelles called Weibel-Palade bodies (WPB)^1^ which owe their unique shape to their main cargo: the pro-haemostatic glycoprotein, von Willebrand factor (VWF).^2^ VWF dimerises in the endoplasmic reticulum and concatemerizes as it passes through the trans Golgi network (TGN) to form long parallel proteinaceous tubules that are then packaged into WPB. Other contents include the pro-inflammatory receptor P-selectin^3^, pro-inflammatory cytokines and agents that control tonicity; thus, exocytosis of WPB is a crucial event important to haemostasis and inflammation.^4^

Regulated secretion of VWF occurs rapidly in response to stimulation with secretagogues released during injury and inflammatory processes.^5,6^ Secreted VWF tubules unfurl to form strings (up to 1 mm long) anchored to the EC surface. These serve as a platform for platelet aggregation and thrombus formation.^7^ This process instigates the primary haemostatic response but is also causally associated with thrombotic diseases such as peripheral vascular disease, myocardial infarction and stroke.^8^ Responsible for one in four deaths,^9^ thrombosis is a leading cause of death world-wide. While current therapy options are numerous, they are complicated by the risk of excess bleeding and cerebral haemorrhage.^10^ As such, there remains a profound medical need for superior treatment strategies.

Circulating levels of VWF are prognostic for cardiovascular disease.^11^ Thus, control of regulated secretion of ultra large VWF multimers is being actively investigated as a therapeutic strategy to reduce the burden of thrombotic diseases. Aptamers and antibodies targeting VWF are currently being tested in the clinic to limit thrombotic pathologies such as thrombotic thrombocytopenic purpura.^12^ We have previously identified cellular machinery that regulates the expulsion of VWF, targeting this process represents an exciting therapeutic approach. ECs recruit actin and myosin during the exocytosis of WPBs to forcibly extrude the ultra large VWF multimers apically (into the blood vessel lumen).^13^ Actomyosin ring compression, aided by septins^14^, is essential for the unfurling of platelet-catching strings across the cell surface.

Myosins are molecular motor proteins that mediate organelle trafficking on actin structures as well as contractile processes on antiparallel actin filaments during muscle contraction, cytokinesis and protein secretion.^15–17^ Conventional class II myosins dimerise as bipolar filaments and by binding to adjacent, oppositely orientated, actin filaments via both head regions the motor proteins exerts force, this is known as the ‘sliding filament hypothesis’.^18^ Class I myosins lack this ability and are referred to as unconventional. They are localised at cell membranes in ruffles, filopodia and the leading edge during migration.^16^ Structurally, class I myosins are composed of an actin-and ATP-binding head domain, a variable neck region and a tail domain.^19^ The ‘neck” (or lever) region contains calmodulin (light chain) binding IQ (isoleucine-glutamine) motif(s) which is thought to act as a regulatory domain, similar to the light chains of class II myosins.^20^ Lastly, class I myosins possess a pleckstrin homology (PH) domain in the tail region that facilitates binding to phosphoinositides.^21^ In some settings, class I myosins transport intracellular vesicles along actin filaments.^22,23^ They have also been shown to tether GLUT4-containing vesicles to actin during exocytosis.^24^ While lung surfactant secreting alveolar type II (ATII) cells utilise actin and class I myosins to aid vesicle compression during lamellar body exocytosis.^25^

A pivotal role of a subset of myosin isoforms in WPB trafficking and VWF secretion have previously been described. WPB are anchored to actin structures in the cell periphery by a tripartite complex of Rab27a, MyRIP and Myosin Va.^26,27^ While non-muscle myosin IIA (NMIIA),^28^ NMIIB^13^ and Myosin Vc^29^ have been implicated in the actomyosin mediated expulsion of VWF. However, the role of class I myosins have not been characterised.

We previously utilised peroxidase proteomics to identify proteins in close proximity to WPB in unstimulated and stimulated conditions.^14^ This powerful approach identified differential proximity of actin-binding motor proteins to the WPB surface in resting ECs and in response to stimuli. Here, we describe the previously undefined function of the class I myosin motor, myosin 1C (Myo1c) and suggest a crucial role in linking the contractile actin ring to the plasma membrane (PM).

## METHODS

### Cell culture

Human Umbilical Vein ECs (HUVEC) (Cat: 12203) and Human Dermal Microvascular ECs (HDMEC) (Cat: 12212) were purchased from PromoCell. ECs were cultured as described elsewhere^30^ with the following exception: HDMEC were cultured using PromoCell Ready-to-use Growth Medium MV (Cat: 22020).

### Immunofluorescent imaging and western blotting

This was performed exactly as described elsewhere.^13^ The commercial suppliers of antibodies used here are as follows. Anti-Myo1c (Abcam, Cat: ab194828), anti-Myo9b antibody (Proteintech, Cat: 12432-1-AP), β-tubulin (Sigma Aldrich, Cat: T4026), VWF (Dako, Cat: A0082 and BioRad Cat; AHP062), LAMP1 (eBioscience, Cat: eBioH4A3) and TGN46 (Bio-Rad, Cat: AHP500GT). Confocal imaging was performed using the Zeiss LSM 800.

### Live cell imaging

GFP-Myo1c was a gift from Martin Bähler (Addgene plasmid # 134832).^31^ GFP-Myo1c-Tail+3IQ, Myo1c-K892A–GFP and Myo1c-R903A–GFP were kind gifts from Michael Ostap.^21^ PH-PLCδ1-GFP/RFP were gifts from Christian Halaszovich.^32^ GFP-PIPK1 gamma 87 was a gift from Pietro De Camilli (Addgene plasmid # 22300).^33^ VWF-GFP was a gift from J. Voorberg and J.A. Van Mourik (Sanquin Research Laboratory, Amsterdam, Netherlands).^34^ P-Selectin lumenal domain mCherry (P.sel.lum.mCherry) was previously cloned in our laboratory.^13^ LifeAct-GFP was a gift from B. Baum (University College London, London, England, UK).^35^ Image J and Circle tool was used to measure Maximum intensity grey values.^36^

### Assessment of target protein inhibition on VWF secretion using NIR fluorescent dot blot

OnTarget plus siRNAs targeting Myo1c (Cat: L-015121-00-0005) were purchased as SMARTpools from Dharmacon (Horizon Discovery). Firefly luciferase targeted siRNA was made by Eurofins Genomics (sequence 5’ cgu-acg-cgg-aau-acu-ucg 3’). Electroporation of HUVEC, VWF secretion assay and near-infrared (NIR)-fluorescent dot blot was performed as described in our previous research.^14^ Phorbol 12Cmyristate 13Cacetate (PMA) (100 ng/mL), histamine (100 μmol/L), thrombin (2.5 U/mL), adrenaline (10 μmol/L) or 3CisobutylC1Cmethyl xanthine (IBMX) (100 μmol/L) were used to stimulate WPB exocytosis. For myosin 1 inhibition, HUVEC were exposed to Pentachloropseudilin (PCLP) (AOBIOUS, Cat: AOB33969) for 16 hours or 30min (5-20 µM) prior to stimulation with secretagogue.

### Data Sharing Statement

R and GraphPad Prism were used to visualise data from our previously published proximity proteomic data set.^14^ Mass Spectrometry data is available via the PRIDE^37^ partner repository with the dataset identifier PXD036983 and 10.6019/PXD036983. For other original data and constructs, please contact.

## RESULTS

### APEX2 proximity proteomics identified differentially enriched myosin isoforms as putative regulators of WPB dynamics

Myosin isoforms have pleotropic functions in secretory vesicle trafficking. They are essential for pre-and post-fusion exocytic processes, including the anchoring of vesicles to peripheral actin, remodelling of cortical actin, stabilisation/linking the fusion pore to the PM and force-driven compression to mediate cargo expulsion^38^. In order to discern which myosin isoforms were of importance in regulated VWF secretion we consulted our publicly available proximity proteomics data set.^14^

A volcano plot was generated to illustrate which myosin isoforms were most upregulated and most significantly enriched proximal to WPBs (Fig 1A). Myosin Va (MYO5A), forms a tripartite complex with Rab27a and MYRIP in order to anchor WPBs to actin structures in the cell periphery.^39^ As expected, Myosin Va was significantly enriched in both unstimulated and stimulated (PMA/HAI) Rab27a-proximity proteomic data sets.^14^ Unexpectedly, the class IX myosin, Myosin 9B (Myo9B) was the most highly enriched and statistically significant myosin isoform proximal to WPBs in both resting and stimulated cells. This unusual myosin motor has a Rho-GAP domain in its tail region.^40^ As Rho activation has previously been implicated in VWF secretion^41,42^ we did not anticipate that Myo9B was a positive regulator of VWF secretion. Indeed, knockdown of Myo9B by siRNA did not affect VWF secretion following exposure to distinct secretagogues (Fig. S1). We speculate that as a focal adhesion resident protein,^43^ Myo9B was detected in our screen due to the close proximity of WPBs to focal adhesions,^28,44^ rather than due to a direct interaction.

**Figure 1:**
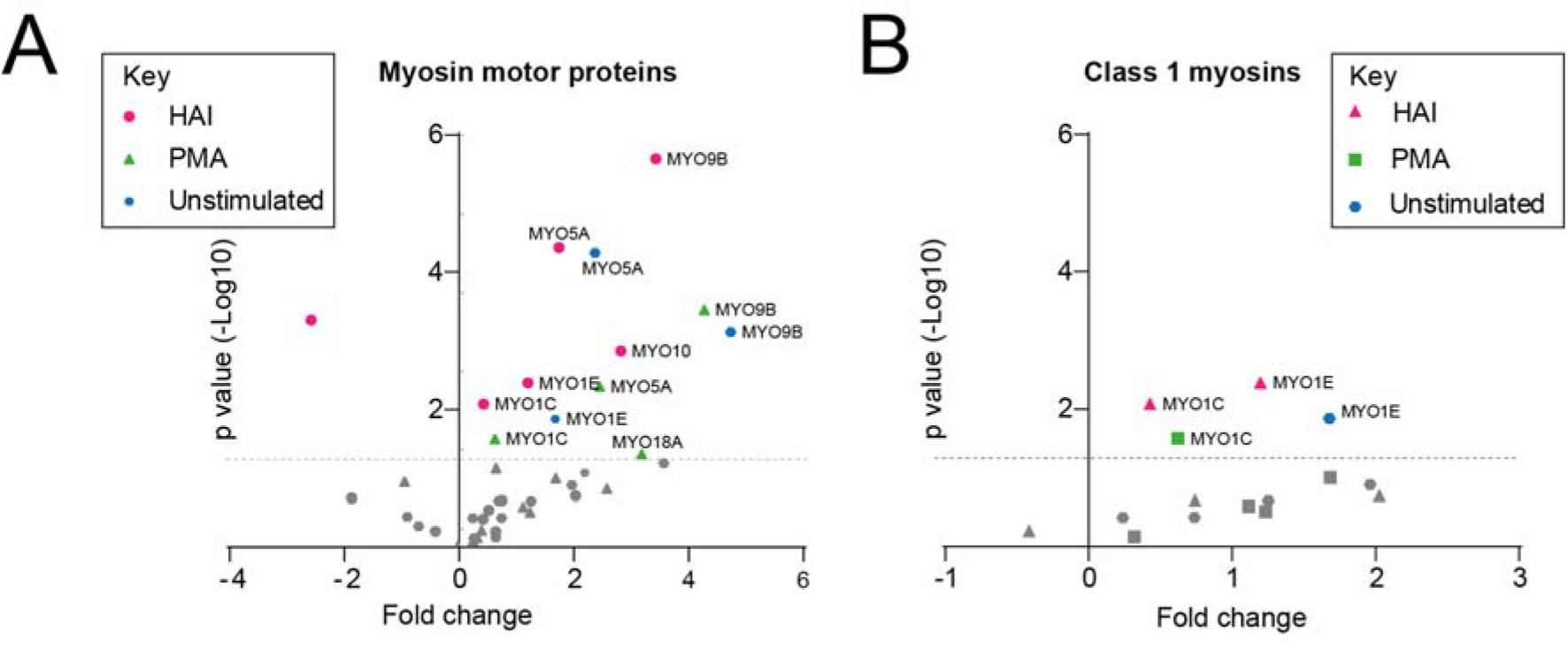
WPB proximal myosin motors. (A) Volcano plot of myosin isoforms in close proximity to WPBs, previously identified by Rab27a targeted APEX2 proximity proteomics. (B) Volcano plot of class I myosin isoforms. Blue - significantly enriched in unstimulated cells. Green – significantly enriched in PMA stimulated cells. Magenta – significantly enriched in HAI stimulated cells. Grey - not statistically significant as compared to mock transfected HUVEC. Paired *t* test.

Next, we chose to investigate the class I myosin family, of which two isoforms were significantly enriched in proximity to WPBs (isoforms C and E) (Fig. 1B). Of these only Myo1c was enriched exclusively following secretagogue stimulation (both PMA and HAI), indicating a potential role in regulated exocytosis.

### Myo1c is recruited during exocytosis

Myo1c has proposed roles in membrane fusion of GLUT4 containing vesicles,^45^ compression of lung surfactant secreting lamellar bodies^25^ and linking actin to the PM during compensatory endocytosis in frog eggs.^46^ Immunofluorescence (IF) studies of unstimulated HUVEC showed Myo1c localised at one side of the cell (Fig 2A, white arrows), likely representing the leading edge of a migrating cell.^47^ Myo1c did not co-localise with VWF/WPBs under these conditions (Fig. 2A: box and inset). Following secretagogue stimulation the WPB fuse with the PM and the rod-shaped WPB collapse as the pH of the organelle shifts from acidic to neutral.^4^ Corroborating our previously reported findings,^14^ Myo1c signal was apparent encapsulating fused WPB (Fig 2A). Utilising the actin poison, cytochalasin E (CCE) with and without stimulus, we noted that Myo1c recruitment is independent of actin (Fig. 2A). As a complimentary approach we utilised live cell imaging of HUVEC transiently expressing GFP-tagged Myo1c^31^. Co-expression with LifeAct-Ruby illustrated colocalization with actin at the leading edge (Fig. 2B). While co-expression with P.sel.lum.mCherry^13^ (a fusion marker which is stored in WPBs and lost upon fusion with the PM) allowed assessment of Myo1c recruitment dynamics during WPB exocytosis. In response to PMA or HAI stimulation, Myo1c-GFP was recruited to WPBs post-fusion and remained present after the fusion marker was lost (Fig. 2C&D).

**Figure 2:**
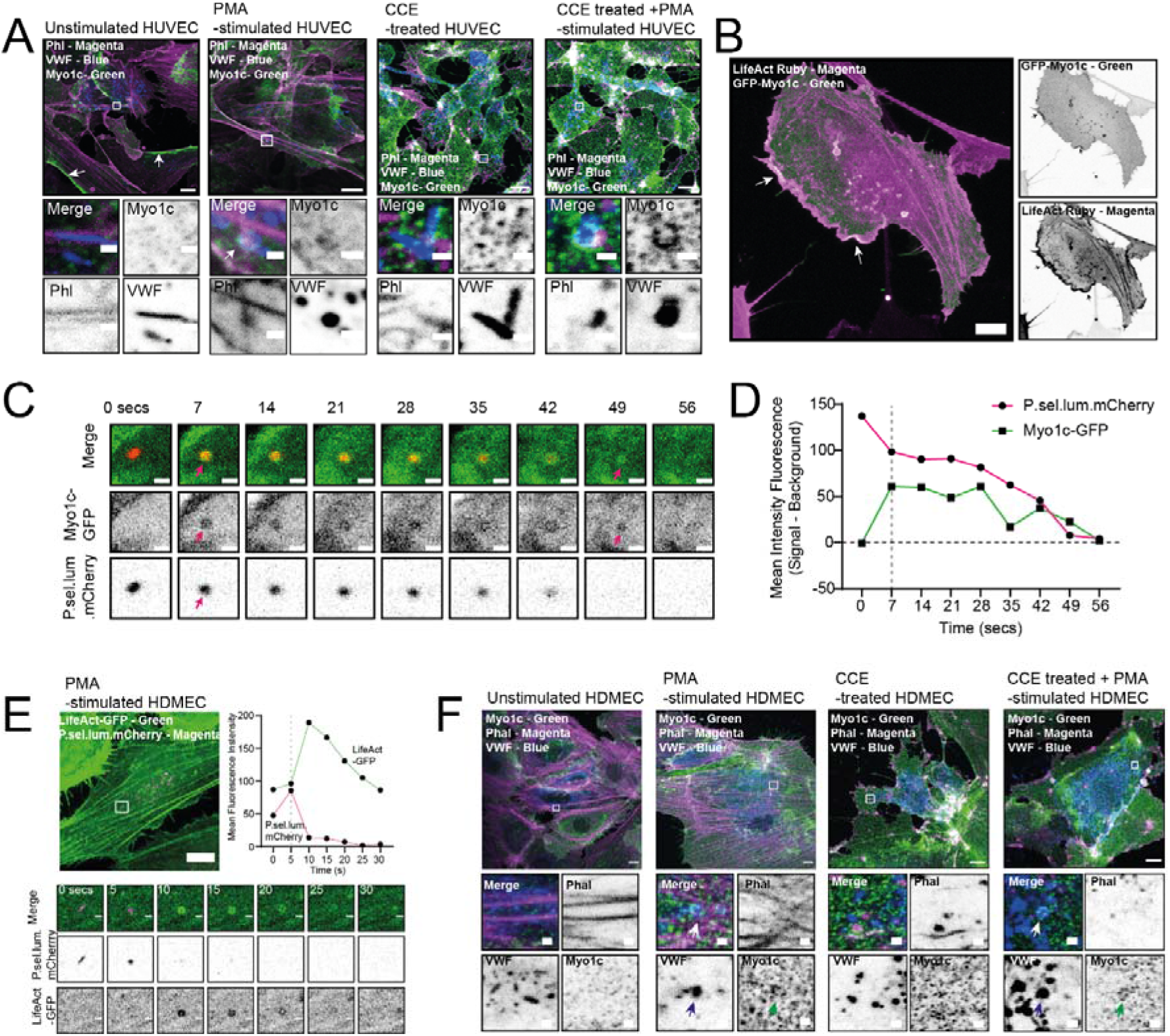
Endothelial cells utilise Myo1c as part of the WPB exocytic machinery. (A) IF localisation of endogenous Myo1c (green), actin (magenta) and VWF (blue) in unstimulated or PMA (100 ng/ml) stimulated HUVEC in the presence and absence of 1 µM of the actin poison cytochalasin E (CCE). Scale bar 10 µm. Inset 1 µm. Arrows indicate polarised localisation of Myo1c. Myo1c is recruited independently of actin but was dependent on stimulation with PMA. (B) Co-expression of GFP-Myo1c (green) and LifeAct-Ruby (magenta) demonstrated col-localisation with actin at the leading edge. (C) GFP-Myo1c encapsulates WPB post-fusion as determined by live cell confocal imaging of HUVEC co-expressing a GFP-Myo1c and the WPB fusion marker P.sel.lum.mCherry. HUVEC were stimulated with 100 ng/ml PMA and imaged live with a confocal laser scanning microscope. Z-stacks at a spacing of 0.5 µm were acquired continuously. Scale bar 1 µm. Arrow indicates point of collapse/fusion of vesicle (D) Recruitment kinetics of GFP-Myo1c (Mean intensity fluorescence). Dashed grey line = fusion/vesicle collapse. (E) Live cell imaging of LifeAct-GFP and P.sel.lum.mCherry expressing microvascular endothelial cells indicated the utility of actin rings to expel VWF following stimulation. Scale bar 10 µm. Inset 1 µm. (F) IF analyses of endogenous Myo1c in HDMEC that were left untreated or stimulated with PMA, CCE or CCE and PMA. White arrows illustrate where GFP-Myo1c is recruited to WPB. Scale bar 10 µm

In order to confirm our results in an alternative EC type we assessed whether microvascular ECs (HDMEC) utilised actin rings during WPB exocytosis. Live cell imaging of HDMEC expressing LifeAct-GFP and P.sel.lum.mCherry confirmed that this phenomenon is not specific to venous ECs from the umbilical vein (Fig. 2E). IF studies demonstrated that HDMEC also recruit Myo1c during VWF secretion. Myo1c did not colocalise with VWF under resting conditions, but when stimulated with PMA, Myo1c could be seen as punctae encircling sphere-like VWF signal (fused WPBs) (Fig. 2F). Once more, this was shown to be independent of actin. This demonstrates HUVEC are a physiologically relevant model for studying Myo1c function and that Myo1c likely has post fusion role. For the remainder of the investigations, we used HUVEC as our model system.

### PIP2 mediated recruitment of Myo1c

Phosphoinositides control membrane traffic and are differentially distributed between cellular compartments.^48^ Myo1c has a PIP2 binding PH domain in its tail region.^21^ Based on research in ATII cells^25^ we anticipated that Myo1c was potentially recruited to fusing WPBs via this region (Fig. 3A). Lipids on the organelle and plasma membrane play a variety of roles in exocytosis in diverse secretory systems.^49^ Phosphatidic acid and PIP2 are likely important in WPB exocytosis as phospholipase D1 (PLD1) has an established role in VWF secretion.^50^ PIP2 sensors (PH-PLCδ1-YFP) and enzymes (PIP5Kγ87) have previously been shown to be recruited to site of WPB fusion to recruit exocytic effector proteins.^51^ We independently confirmed that GFP-PIPK1γ87 (Fig. 3B) and PH-PLCδ1-GFP (Fig. 3C) were present at sites of WPB fusion following collapse of the organelle. The kinetics mirrored that of Myo1c-GFP recruitment-post-fusion. Interestingly, we on occasion noted the presence of GFP-positive vacuoles in GFP-PIPK1γ87 expressing cells that were coated in Myo1c, actin and septin 7 (another lipid binding protein associated with WPB exocytosis) (Fig. S2). These data indicate the presence of PIP2 on the plasma membrane drives protein recruitment.

**Figure 3:**
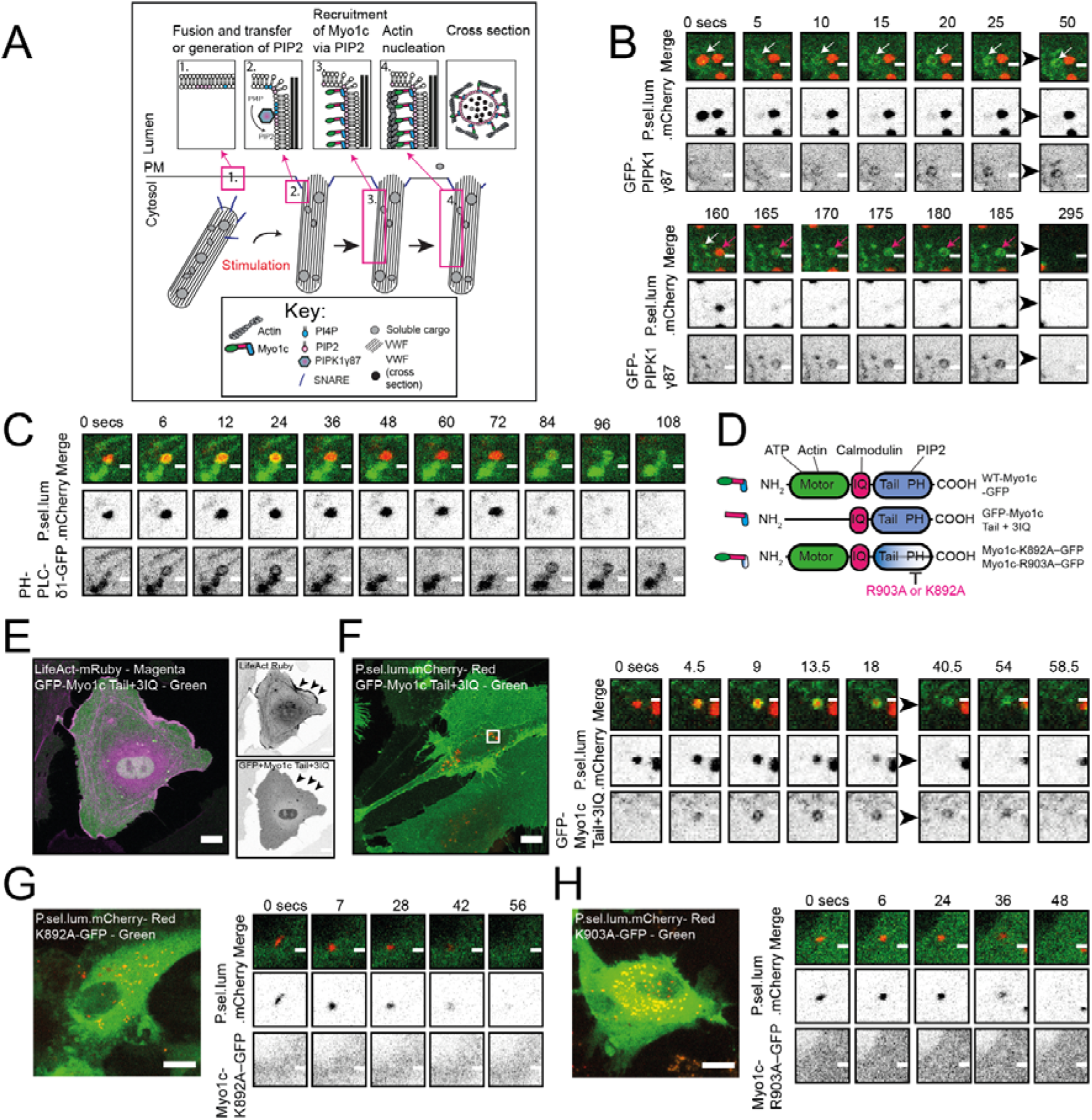
The PH domain of Myo1c is required for its recruitment during WPB exocytosis. (A) A schematic of the proposed spatiotemporal dynamics of Myo1c recruitment during WPB exocytosis. (B) Live cell imaging of the GFP-PIPK1γ87 and P.sel.lum.mCherry in secretagogue (HAI) stimulated HUVEC illustrated its recruitment post-fusion. Two exocytic events are seen here. Scale bars are 1 µm. White and magenta arrows indicate independent fusion events (C) The PIP2 sensor PH-PLCδ1-GFP was also recruited post fusion. Scale bars are 1 µm (D) A schematic of the Myo1c structural domains and location of truncation or mutations. (E) HUVEC co-expressing LifeAct-Ruby and GFP-Myo1c Tail+3IQ indicated the importance of the myosin head domain for interacting with actin. Scale bars are 10 µm (F) GFP-Myo1c Tail+3IQ was recruited to WPBs post-fusion (G&H) GFP-tagged Myo1c fusion proteins harbouring mutations in their PH domain (K892A/R903A) were not recruited to WPBs during exocytosis. F, G & H Scale bars are 10 µm. Inset scale bars are 1 µm.

To investigate the mechanism of its recruitment we used truncated and point mutants of Myo1c (Fig. 3D). By co-expressing the GFP-tagged neck and tail domain of Myo1c (GFP-Myo1c-Tail + 3IQ) together with LifeAct-Ruby we demonstrate that the N-terminus is necessary for appropriate Myo1c targeting to actin at the leading edge (Fig. 3E and inset). Consistent with the hypothesis that Myo1c is recruited to fusing WPB via its tail-resident PH domain; GFP-Myo1c-Tail + 3IQ localised to fused WPBs following secretagogue stimulation (Fig. 3F). We used a GFP-tagged Myo1c fusion proteins harbouring point mutations in the phosphoinositide-binding PH domain (K892A/R903A) to show that the interaction with PIP2 is necessary for recruitment to WPB at fusion (Fig. 3G&H). Unlike the wild type construct Myo1c-Tail + 3IQ, Myo1c-K892A–GFP and Myo1c-R903A–GFP localise to the cytosol and are not recruited to WPBs post-fusion. Taken together, these data indicate that Myo1c is recruited to WPB post-fusion via its PH domain.

### The effect of Myosin-1 inhibition of exocytic actin ring dynamics

To investigate the role of the motor (head) domain we utilised the pan-myosin I inhibitor, pentacholoropseudilin (PCLP)^52^. PCLP is a potent allosteric inhibitor of myosin ATPase which shows selectivity for class I myosins at low doses (IC50 ∼ 1-10 µM). However, at higher doses, other myosin classes (e.g. NMIIB IC50 ∼ 90 µM)^52^ are affected. Here, pre-exposure to 10-20 µM PCLP for 16 hours resulted in an obvious trafficking defect whereby total levels of mature VWF were decreased in a dose dependant fashion and the ratio of pro-VWF to mature VWF increased at the highest dose (Fig. S3A-D). VWF secretion (Fig. S3E & F) and string formation (Fig. S3G-I) was almost completely inhibited. Moreover, IF of LAMP1 (Fig. S3J) and TGN26 (Fig. S3K) illustrated VWF positive lysosomes and a gross defect in morphology of the TGN. This demonstrates that class I myosins play a role in different stages of the secretory pathway.

Acute PCLP incubation (30 minutes) allows post fusion analysis of the role of myosin 1 in WPB exocytosis. We used live cell imaging of LifeAct-GFP and P.sel.lum.mCherry (Fig. 4A). Live cell imaging demonstrated that Myosin 1 inhibition had no effect on the percentage of WPB fusion events that recruited an actin ring (Fig. 4C). This indicated that in this system, Myo1c is not required for actin polymerisation. However, an increased proportion of persisting (>60s) actin coats/rings in PCLP treated cells indicated a potential role in augmenting compression potentially through lack of actin coat organisation cause by improper actin-membrane linkage (Fig. 4D).

**Figure 4:**
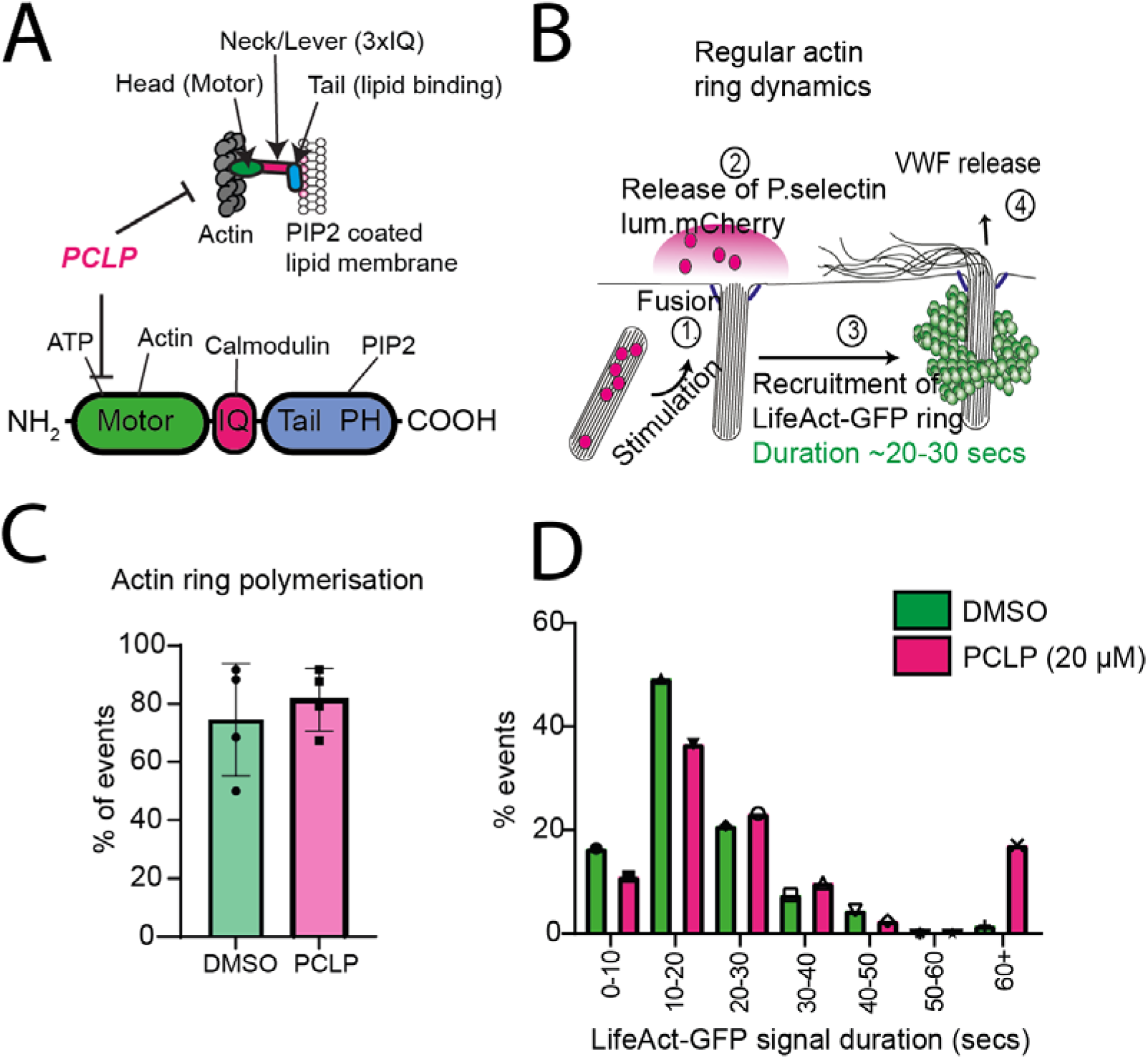
Acute inhibition of class I myosins changes actin ring dynamics. (A) Schematic of Myo1c domains and mechanism of inhibition by PCLP. (B) Schematic of live cell imaging approach to study actin dynamics during WPB exocytosis. (C) HUVEC co-expressing LifeAct-GFP and P.sel.lum.mCherry were stimulated with PMA (100 ng/mL) and image continuously (0.5 µM Z stacks) for 10 minutes. The percentage of WPB fusion events that recruited an actin ring were unchanged in DMSO and PCLP (20 µM) treated cells. [14 (DMSO) and 16 (PCLP) cells analysed across 4 experiments, mean ± SEM] (D) The lifetime (secs) of LifeAct-GFP signal at fusion sites was quantified in DMSO (67 events) and PCLP (82 events) treated HUVEC. The distribution of frequency of events is presented here (across n=4).

In rare events (3-4%), we saw excessive actin polymerisation during ring formation which resulted in actin comet tail formation (Fig. S4A & B), reminiscent of those noted during bacterial and viral infection.^53^ This indicates that Myo1c may also constrain actin polymerisation at the membrane to allow correctly polarised force production. Although the relative incidence of these events was low, they suggest that during WPB exocytosis, Myo1c organises and links actin to the WPB allowing directional compressive force during VWF secretion. We noted that cytosolic comet tails were commonly observed in HUVEC transiently overexpressing GFP-PIP5Kγ87 (Fig. S4C&D). Co-expression of ARP3-mCherry indicated that a proportion of these comet tails were generated by the ARP2/3 complex (Fig. S4E).

Taken together, our data indicates that the myosin ATPase domain is required for Myo1c exocytic function, consistent with a role in linking the actin ring to the WPB surface. We hypothesise that without proper linkage to the WPB membrane, excessive PIP2 mediated actin polymerisation results in actin rings with disorganised mechanics resulting in mistargeted release of VWF.

### Myo1 influences WPB exocytosis

To assess the functional role of Myo1c on regulated VWF secretion we depleted the endogenous pool of Myo1c. To limit off target effects^54^ we utilised a pool of the four oligonucleotides. This resulted in knock down efficiencies of 71-88% across three independent experiments (Fig 5A). In agreement with a role in WPB exocytosis, Myo1c knock down reduced VWF secretion in response to PMA (Fig. 5B) and a more physiological regulator of VWF secretion, thrombin (Fig. 5C).

**Figure 5:**
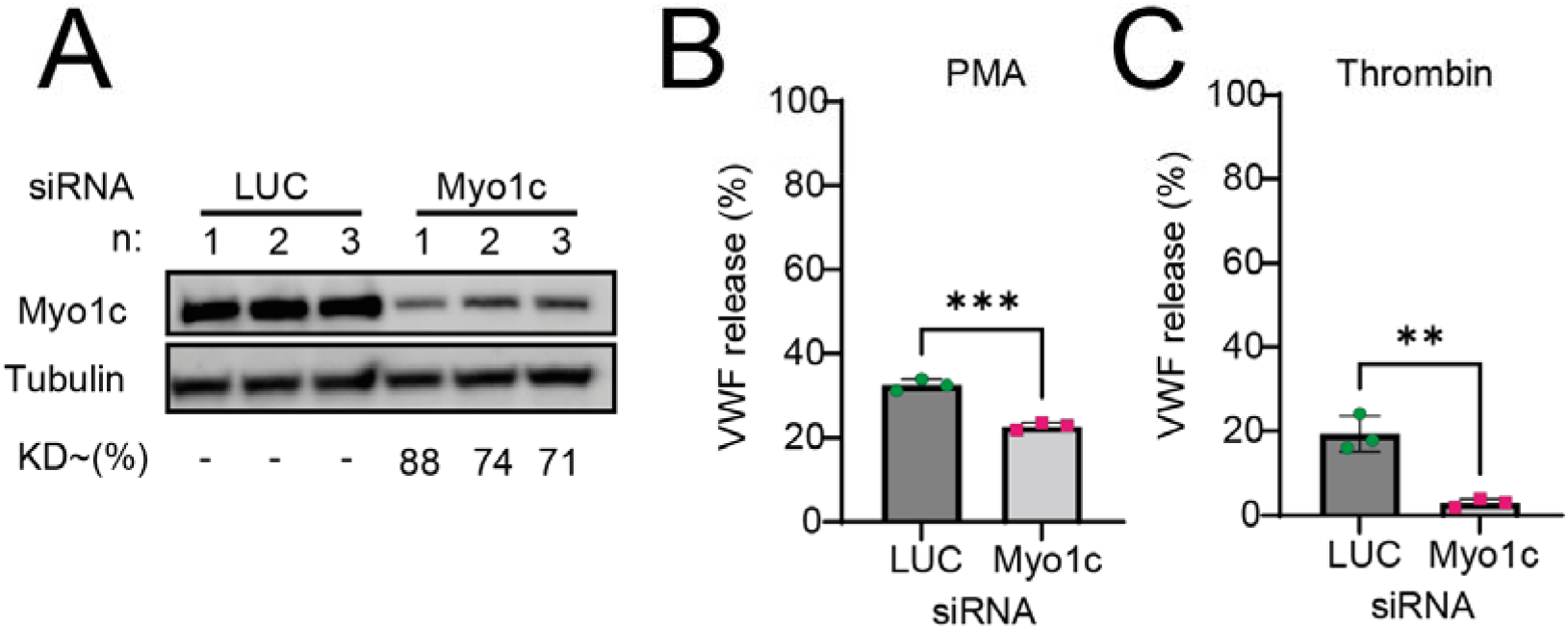
Myo1c is required for the efficient expulsion of VWF from HUVEC. A pool of four independent Myo1c-targeting siRNA oligonucleotides was used to deplete Myo1c. (A) Immunoblotting was used to confirm target depletion. VWF secretion in response to (B) PMA (100 ng/mL) and (C) thrombin (2.5 U/mL) was next assessed by NIR dot blot. n=3. Students *t* test. ***P<0.005 **P<0.01. KD: Knock down (KD) of target protein concentration as normalised to tubulin.

Because prolonged exposure of PCLP resulted in a VWF trafficking defect, we next assessed the effect of four different siRNA oligonucleotides on VWF secretion and total VWF levels by western blotting. Western blotting determined that each oligonucleotide significantly reduced Myo1c protein levels but with differing degrees of efficiency (45-84%) (Fig. S5A&B). Convincingly, VWF secretion in response to HAI correlated with levels of endogenous Myo1c (Fig. S5C). Levels of mature and pro-VWF were unchanged by Myo1c knockdown indicating that Myo1c is not required for WPB biogenesis. The broader effect of PCLP likely reflect effects on other class I myosins (Fig. S5D-G).

## DISCUSSION

We previously described the presence of a contractile actomyosin ring that is recruited to the WPB surface following fusion with the PM and aiding efficient VWF secretion.^13^ This represents an unexploited therapeutic target for the prevention of thrombotic pathologies. Actomyosin mediated expulsion of VWF requires upstream protein kinase C α (PKCα)^55^ and p21 activated kinase 2 (PAK2)^14^ signalling, Spire1 mediated actin nucleation^56^, zyxin and α-actinin mediated organisation^57^ and controlled compression (septin^14^ and non-muscle myosin isoforms^13,28^). However, the mechanism by which the actomyosin coat/ring is attached to the vesicle membrane and how this influences exocytosis is unclear. Here, we demonstrate that Myo1c is recruited to the membrane of fused WPBs by its PH domain, in an actin independent fashion. Perturbation of Myo1c function through siRNA mediated depletion reduced VWF secretion in human ECs and is thus of fundamental physiological importance to haemostasis and thrombosis.

Although other class I myosins such as Myo1e^58^ have reported roles in other secretory systems, we have focused here on delineating a role for Myo1c. In addition to actin-membrane tethering, Myo1c has also been reported to aid insulin stimulated PM-fusion of GLUT4 containing vesicles, where it is recruited prior to vesicle fusion.^22,59^ In contrast, we observed that GFP-tagged Myo1c is only recruited to the WPB surface post-fusion, similar to that reported by Kittelberger and colleagues.^25^ We therefore exclude the possibility that Myo1c is acting as an organelle transporter. Instead, we propose a similar role to that seen in surfactant exocytosis by ATII cells^25^ and cortical granules in Xenopus eggs^46^ whereby Myo1c links the actin coat to the vesicle membrane. To fulfil this role, Myo1c would need to have a tight interaction with both actin and the WPB membrane surface. Concordantly, the PH domain of Myo1c has previously been shown to bind PIP2, which we and others^51^ demonstrate are recruited to the WPB post-fusion. Furthermore, we show here that the recruitment of Myo1c is dependent on this region. Next, inhibition of the myosin head ATPase using PCLP perturbed actin ring dynamics, although it did not reduce the proportion of fusion events that recruited actin. An increased incidence of longer lasting actin rings (+60s) was observed. These data closely resemble the defect in actin ring contraction observed when NMII isoforms are inhibited using blebbistatin.^13^ In rare cases we observed excessive and misdirected actin polymerisation in the form of a comet tail, such as seen in cortical granules when Myo1c was inhibited using a dominant negative construct.^51^ Similar phenomena has previously been reported in ATII cells^60^ and in Xenopus eggs when Myo1c was inhibited through expression of a dominant negative mutant (Tail+3IQ).^46^ Likewise, during actin-mediated phagocytosis, Myo1e and Myo1f double knock-out macrophages display excessive actin polymerisation at the phagocytic cup which results in prolonged or delayed phagocytosis.^61^ We speculate that under normal condition Spire1 controls actin ring recruitment^56^ while these rare comet tails are caused by excess PIP2 mediating N-WASP/ARP2/3 actin polymerisation.

Here, an acute PCLP exposure time of 30 minutes was needed to assess actin ring dynamics during WPB exocytosis. Longer incubation times had drastic effects on protein trafficking. Sixteen hours incubation with PCLP resulted in disruption of the TGN, a near complete abrogation of regulated VWF secretion, ineffective VWF biogenesis as well as the clear observation of VWF signal in LAMP1 positive lysosomes. These may reflect defects at the Golgi^62^ or by regulating autophagosome lysosome fusion^63^ with subsequent effects on WPB turnover, lysosomal co-localisation and degradation of VWF. Importantly, we did not observe this phenotype in Myo1c knock down cells and therefore hypothesise that this trafficking defect is caused by the indiscriminate inhibitory action of PCLP on other class 1 myosins.^52^ Notably, Myo1b promotes the formation of tubules and carriers at remodelling TGN membranes^64^ and has a role in secretory granule biogenesis in pancreatic Beta cells^65^ and neuroendocrine cells.^66^A specific role in WPB biogenesis is therefore also a possibility.

As a functional read out, we assessed the effect of siRNA mediated depletion of Myo1c on VWF secretion in response to PMA, HAI or thrombin. These secretagogues were specifically chosen as they are known to stimulate increases in both cytosolic cAMP and Ca^2+^.This was important as cAMP is thought to be required for ring recruitment^28,55,57^ and Ca^2+^ required to stabilise the Myo1 lever arm when in action.^20,67^ In agreement with our detailed image analyses, Myo1c depletion reduced VWF secretion suggesting an active role in actomyosin mediated expulsion of VWF possibly through facilitating membrane tension and likely in concerted action with class II myosins.^13,28^

Here we reported that siRNA depletion of Myo1c modestly reduced VWF secretion in response to PMA but a greater effect was observed when HUVEC were stimulated with thrombin. This may be due to distinct signalling pathways used by these secretagogues. PMA is known to induce VWF secretion activation of PKC while thrombin is known to stimulated Ca^2+^ and calmodulin signalling.^68^ Myo1c is a calcium sensitive and calmodulin binding domain-containing motor protein and as such this may explain our findings. Alternatively, phospho-proteomic analysis of HUVEC stimulated with thrombin previously identified that Myo1c is rapidly phosphorylated at serine 29.^69^ It is therefore plausible that post-translational modification of Myo1c may be important in governing its function.

Finally, as a hypothesis of the spatiotemporal recruitment of these molecules we present a putative working model (Fig. 6). Enriched PIP2 concentrations occur at the site of WPB fusion either as a result of membrane mixing and/or through PIPK1γ87 mediated production from its precursor PI4P. PIP2 at the WPB surface then leads to the rapid recruitment of Myo1c via its lipid binding PH domain. De novo Spire1 mediated actin nucleation^56^ follows (or occurs simultaneously) with the Myo1c motor domain binding to the resulting actin coat/ring. We suggest that NMII isoforms are recruited after actin, such as seen in Xenopus eggs^70^, lamellar bodies,^71^ rodent^72^ and Drosophila salivary granules.^73^ Activation of these isoforms then leads to vesicle compression and expulsion of VWF.

**Figure 6:**
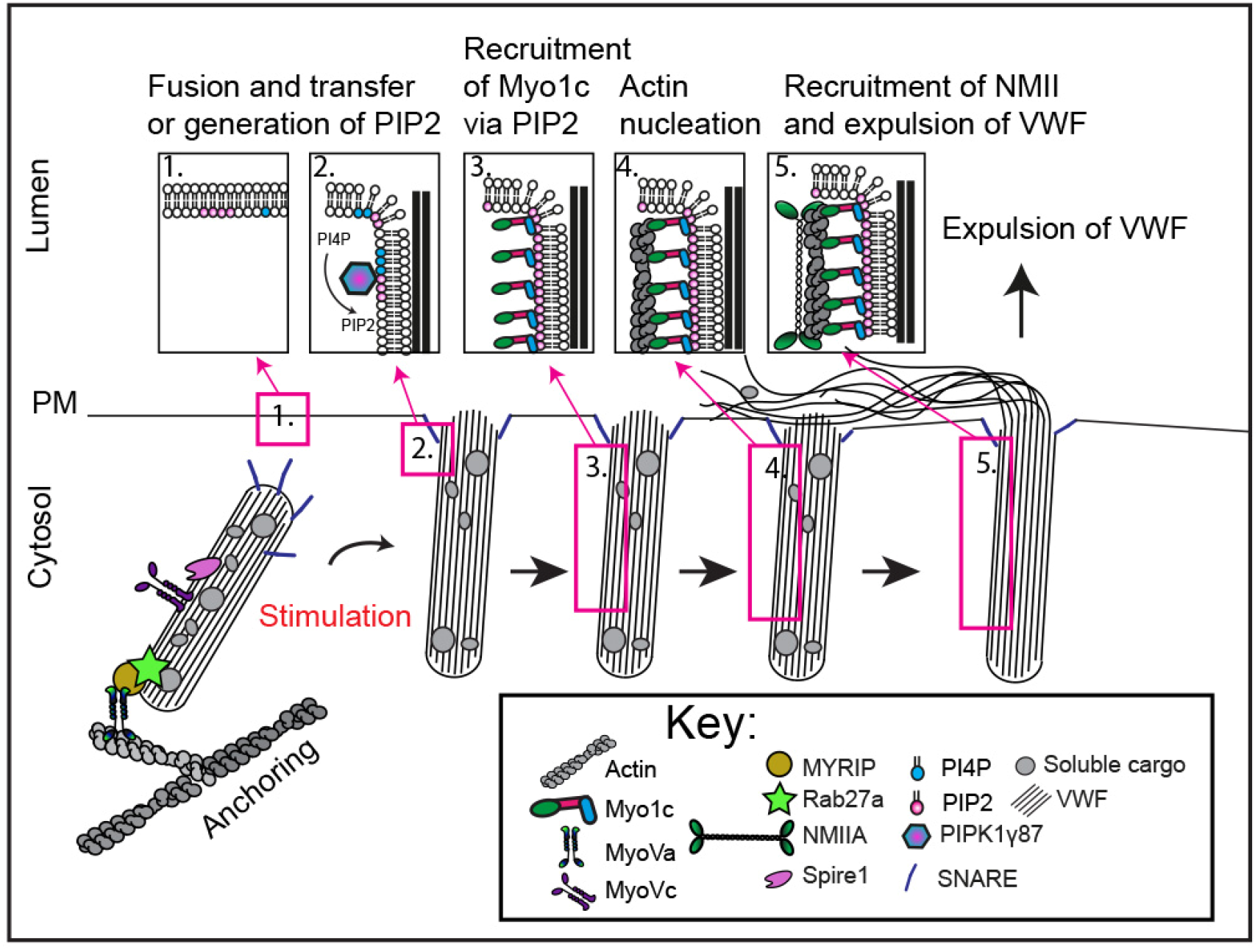
A working model for actomyosin mediated expulsion of VWF. Under resting conditions WPB are anchored to peripheral actin structure via Rab27a/MyRIP/MyoVa. (1) Following stimulation WPB are trafficked to the plasma membrane where they will fuse. (2) An enrichment of PIP2 is present at the site of fusion, possible also on the WPB surface. This could be through membrane mixing or through the catalytic activity of PIPK1γ87. (3) Myo1c is recruited to the WPB via its PIP2 binding PH domain. (4) Spire1 mediated de novo actin nucleation could occur after Myo1c recruitment or simultaneously. (5) Recruitment of NMII isoforms likely happens after actin polymerisation. Activation of NMII allows for the expulsion of VWF into the blood vessel lumen.

## CONCLUSION

Overall, these data provide the first evidence of class I myosins participating in VWF secretion from ECs. And to our knowledge this is the first description of a role for Myo1c in the field of thrombosis and haemostasis. As such these data aid our fundamental understanding of the molecular mechanisms governing primary haemostasis.

## ACKNOWLEDGMENTS

We acknowledge the CMR Advanced Bio-Imaging Facility of QMUL for the use, help and advice with microscopy. We thank Dr. Chris Stefan (University College London) for his critical appraisal of this manuscript.

## SUPPLEMENTAL

### Supplemental figure 1

**Figure S1:**
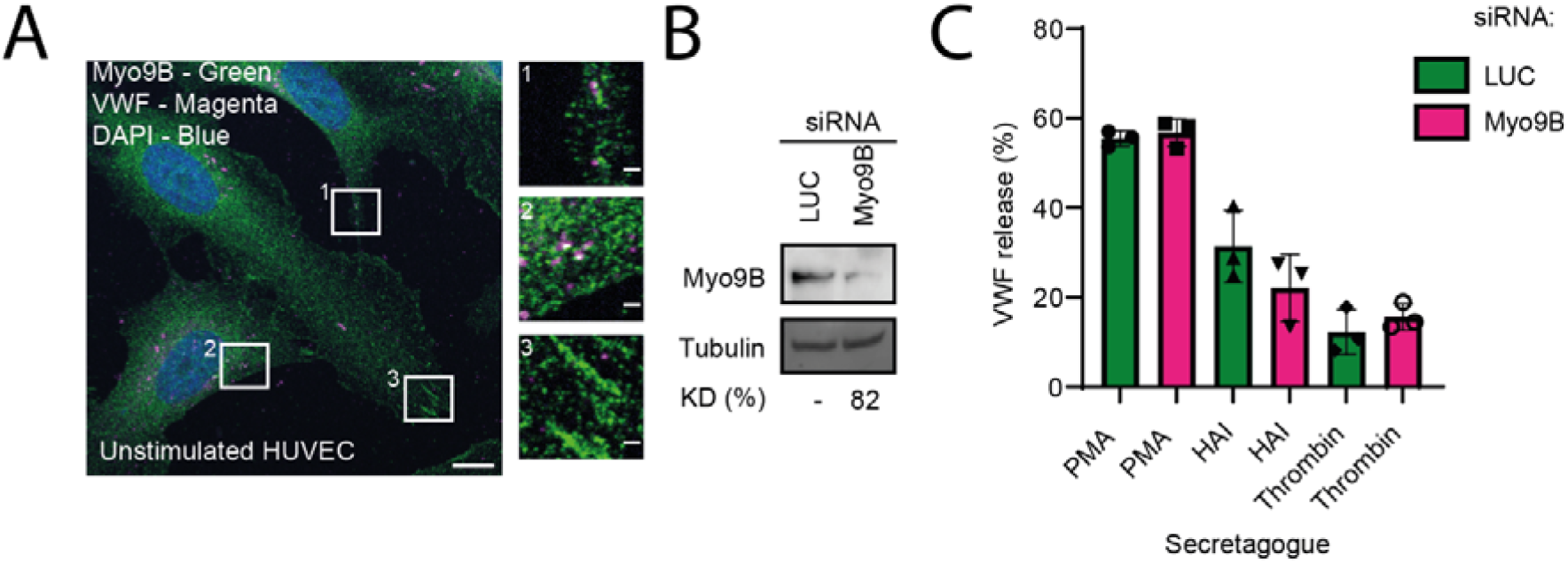
Myo9b is not required for VWF secretion. (A) Unstimulated HUVEC were fixed and subject to immunofluorescence analysis to localised Myo9B (green) in relation to VWF (magenta). Myo9B staining was present in the cytoplasm and as patch like staining reminiscent of focal adhesions (Inset 1 and 3). In some cases, VWF could be seen to co-localise with Myo9B puncta (Inset 2) Scale bar 1µm. (B) Western blotting of tubulin and Myo9b following two rounds of electroporation of 300 pM luciferase (LUC) and Myo9B targeting siRNA. Representative blot. KD= knock down efficiency (C) VWF secretion in response to PMA, HAI and thrombin was assessed by NIR dot blot n=3.

**Figure S2:**
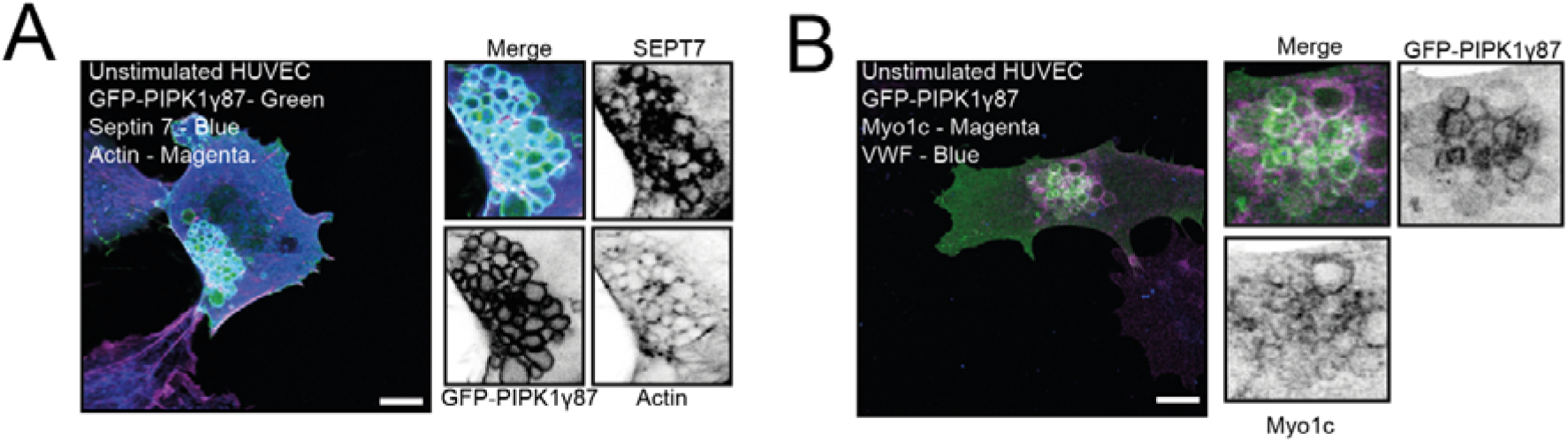
GFP-PIPK1_γ_87 positive vacuoles co-localise with Septin 7, actin and Myo1c. (A) Unstimulated HUVEC transiently expressing GFP-PIPK1γ87 (green) were fixed and probed for septin-7 (septin 7) by immunofluorescence and co-stained with phalloidin to visualise actin (magenta). (B) Myo1c (Magenta) also co-localised with GFP-PIPK1γ87 positive vacuoles. Scale bar 10 µm

**Figure S3:**
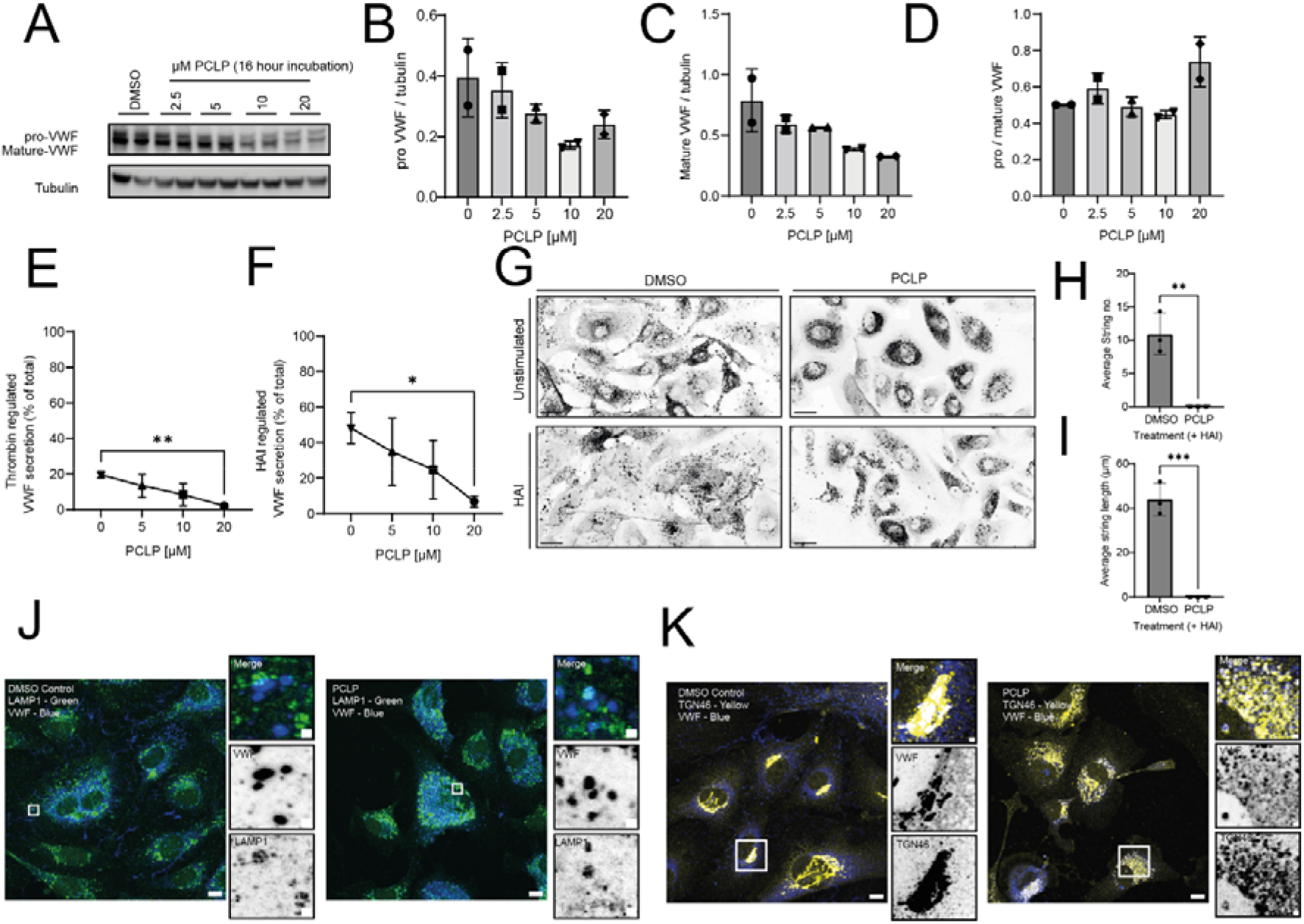
HUVEC exposed to PCLP for 16 hours exhibit a VWF trafficking defect. HUVEC were expose to DMSO or a range of concentration of PCLP and incubated overnight (16 hours) at 37°C 5% CO^2^. (A) Immunoblotting of the resulting lysates displayed changes in pro-and mature-VWF in relation to tubulin. Densitometry indicated a dose dependent decrease in (B) mature-and (C) pro-VWF levels alongside an increase in the (D) ratio of pro/mature VWF at the highest con centration of 20 µM. (E) 16-hour incubation with PCLP resulted in inhibition of regulated secretion of VWF in response to thrombin and (F) HAI. (G) HUVEC pre-incubated with DMSO or PCLP for 16 hours were stimulated with HAI for 10 minutes before application of 5 dyne/cm^2^ shear stress. VWF strings were visualised by immunofluorescence. (H) The number and (I) length of VWF strings secreted under flow in response to HAI in the presence or absence of DMSO or PCLP (n=3). (J) IF analyses using anti-LAMP1 (green) and anti-VWF (blue) antibodies indicated numerous VWF positive lysosomes in PCLP treated cells. (I) IF analyses using anti-TGN46 (yellow) and anti-VWF (blue) antibodies indicated a gross defect in TGN morphology (fragmented and swollen) in PCLP treated cells. Scale bars 10 µm. Inset scale bars 1 µm.

**Figure S4:**
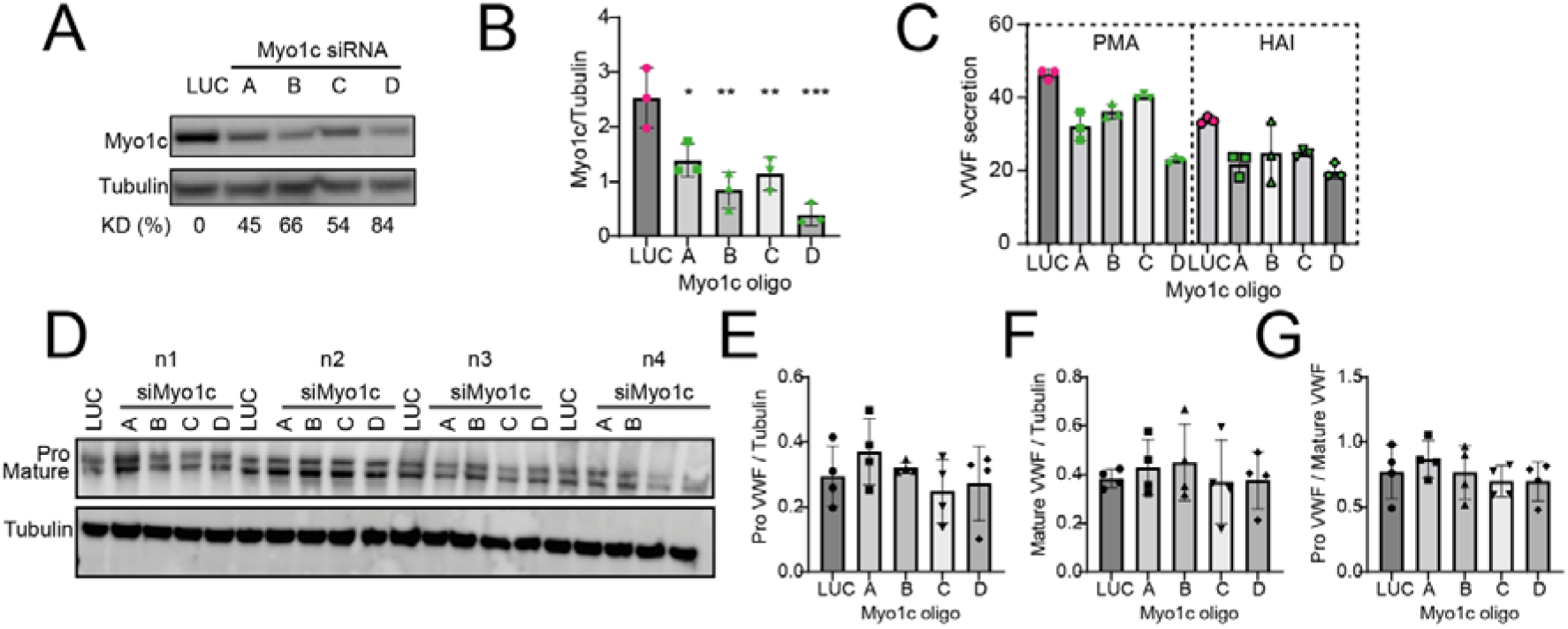
Actin comet tails can occur during WPB exocytosis. (A&B) In rare events (3% mean of 4 independent experiments) excessive actin polymerisation was observed resembling an actin comet tail. Mean ± SEM Scale bar 1µm. (C&D) Cytosolic actin comet tails (magenta) are also seen in HUVEC over-expressing PIP5Kγ87-GFP (green). Scale bar 10µM. Inset 1µM. (E) Presence of ARP3 mCherry on comet tails indicated their polymerisation mediated partly by ARP2/3 complex. Scale bar 10µM. Inset 1µm.

**Figure S5:**
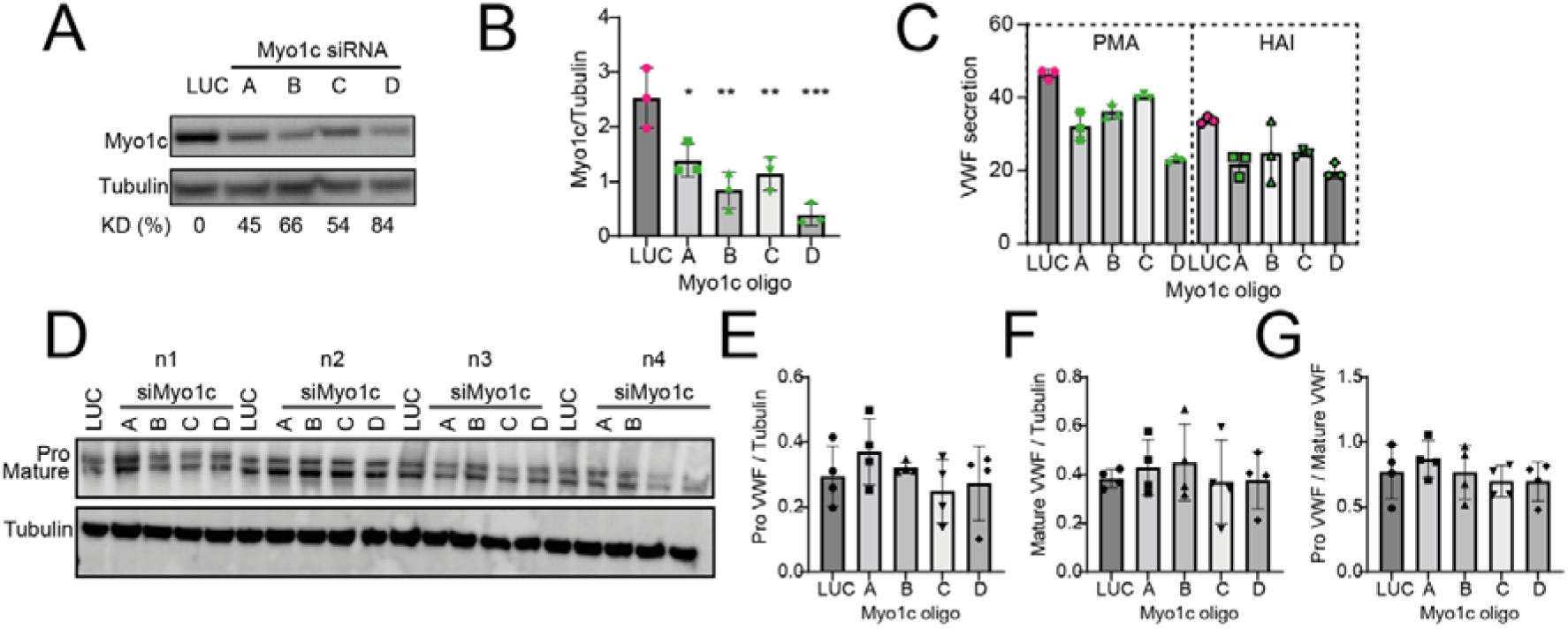
siRNA depletion of Myo1c reduces VWF secretion but does not affect total VWF levels or maturation. Myo1c was depleted using four independent siRNA oligonucleotides targeted Myo1c mRNA (300 picomole). (A) Target protein knock-down was determined using (A) western blotting and (B) densitometry. One way ANOVA with Tukey’s post hoc comparison (C) HUVEC from the same experiments were exposed to PMA or HAI for 30 minutes and levels of VWF secreted into the culture media assessed by NIR fluorescent dot blot. VWF secretion correlated with endogenous levels of Myo1c. Representative VWF secretion values are shown. (D) Immunoblotting of control and Myo1c knock down lysates. Total levels of (E) pro-VWF and (F) mature VWF were not changed following Myo1c KD. (G) the ratio of pro to mature VWF was unchanged following Myo1c indicating that VWF maturation was unaffected. (n=4).

